# Interstitial spaces are continuous across tissue and organ boundaries in humans

**DOI:** 10.1101/2020.08.07.239806

**Authors:** Odise Cenaj, Douglas H. R. Allison, R Imam, Briana Zeck, Lilly M. Drohan, Luis Chiriboga, Jessica Llewellyn, Cheng Z Liu, Young Nyun Park, Rebecca G. Wells, Neil D. Theise

**Affiliations:** Department of Pathology, New York University Grossman School of Medicine, New York NY USA; Divison of Gastroenterology, Department of Medicine, Perelman School of Medicine at the University of Pennsylvania, Philadelphia PA United States; Department of Pathology, Department of Pathology, Yonsei University College of Medicine, Seoul South Korea; Department of Bioengineering, School of Engineering and Applied Sciences, The University of Pennsylvania, Philadelphia, PA, United States; Department of Pathology and Laboratory Medicine, Perelman School of Medicine at the University of Pennsylvania, Philadelphia, PA, United States; Center for Engineering MechanoBiology, The University of Pennsylvania, Philadelphia, PA, United States

## Abstract

Bodies have “reticular networks” comprising collagens, elastin, glycosaminoglycans, and other extracellular matrix components, that are continuous within and around all organs. Fibrous tissue coverings of nerves and blood vessels create structural continuity beyond organ boundaries. We recently described fluid flow through such human fibrous tissues. It remains unclear whether these interstitial spaces are continuous through the body or are discontinuous, confined within individual organs. We investigated IS continuity using two approaches. Non-biological particles (tattoo pigment, colloidal silver) were tracked within colon and skin interstitial spaces and into adjacent fascia. We also exploited hyaluronic acid, a macromolecular component of interstitial spaces. Both techniques demonstrate continuity of interstitial spaces within and across organ boundaries, including within perineurium and vascular adventitia traversing organs and the spaces between them. We suggest a body-wide network of fluid-filled interstitial spaces with significant implications for molecular signaling, cell trafficking, and the spread of malignant and infectious disease.

## Introduction

The work of Franklin Mall over a century ago^1,2^ as well as modern day decellularization techniques^3^ demonstrate that there are “reticular networks” made up of collagens, elastin, glycosaminoglycans, and other extracellular matrix (ECM) components surrounding, within, and between organs. These networks have biological and mechanical roles in defining the architecture and physiology of organs and, as a result, are now used as scaffolding for the creation of customized organ grafts for regenerative medicine.^4^ Multiorgan decellularization has further confirmed that ECM networks extend beyond the confines of single organs to involve neighboring structures, including thoracic (heart), abdominal (liver, gut, kidneys) and pelvic (uterus, prostate, urinary bladder) organs with their vasculature and surrounding fibrous, adventitial sheaths, creating structural continuity across organ boundaries.^5^ More recently, decellularization of entire fetal sheep shows that the connective tissue network is continuous throughout the body and that the connective tissue of nerves creates structural continuity between the nervous system and other tissues.^3^ Dissection of human bodies likewise demonstrates continuity across large, multiorgan regions of the body, including the entirety of the dermis and the fascia of diverse organs and organ systems.^6,7,8,9^

Clinical disciplines including osteopathy have suggested that these connective tissue networks contain fluid and represent a body-wide communications network, akin to interstitial spaces, although this lacks detailed microscopic confirmation. Interstitial spaces in which nutrient and waste exchange take place have historically been recognized at two scales: intercellular spaces (≤1 micron) and pericapillary spaces (~ 10 microns).^10^ We recently described fluid flow through large interstitial spaces of the human extrahepatic bile duct submucosa and the human dermis, 50-70 μm below the epithelial surface.^11,12^ We further showed that other fibrous tissues, including the submucosae of all other visceral organs and the subcutaneous fascia, are structurally similar, and hypothesized that they likewise support fluid flow. In all of these tissues, the spaces were defined by a network of collagen bundles 20-70 μm in diameter. Many of the collagen bundles were lined by spindle-shaped cells that co-expressed vimentin and CD34, but were devoid of endothelial ultrastructural features and were thus considered fibroblast-like cells. In this context, we refer to them as “interstitial lining cells.” In vivo endomicroscopy has shown that musculoskeletal fascia includes similar large-caliber fluid-filled spaces.^13^ It remains unclear, however, whether these interstitial compartments are continuous through the body or represent discontinuous fluid-filled channels confined within individual organs. A limited demonstration of such intra-tissue continuity is found in the work of Mall, who reported that the interstitial spaces of liver portal tract stroma are continuous between peri-arterial, peri-venular, and peri-ductal compartments (the “space of Mall”),^1^ but these studies were only at the local, intrahepatic level.

We investigated the question of interstitial space continuity using two orthogonal approaches. The first was to study movement of non-biological particles (tattoo pigment and colloidal silver) across tissue compartments within colon and skin and into adjacent fascia. The second exploited hyaluronic acid (HA), a macromolecular component of the smallest interstitial spaces (i.e. between cells and around capillaries) as well as the larger fibroconnective tissue spaces we recently identified.^10,14–16^ We demonstrate continuity across organ boundaries and between spaces in all fibrous tissues studied, including the perineurium and vascular adventitia within them. We suggest that there is a broad and interconnected network of interstitial fluid-filled channels throughout the body, including the structural coverings of nerves and vessels, and that this has significant implications for molecular signaling, cell trafficking, and the spread of malignant and infectious disease.

## Results

### Tattoo pigment and colloidal silver are found at a significant distance from original sites of application

Skin samples with tattoos injected into dermis for cosmetic reasons were examined for the presence of pigment particles distant from the dermis. In 3 samples obtained from different patients, particles were identified in papillary and reticular dermis and subcutaneous fascia (**Fig. 1a**). The particles were localized both intracellularly, within the cytoplasm of macrophages and interstitial lining cells, and extracellularly, within interstitial spaces between collagen bundles of the collagenous network of the dermis and subcutaneous fascia (**Fig. 1b,c)**. Silver particles were observed in similar locations in two samples from a patient who developed argyria after topical application of colloidal silver (**Fig. 2a-f**). Silver particles were also identified in the adnexa, perivascular adventitia and perineurium in the dermis (**Fig. 2g-l**).

**Fig. 1.**
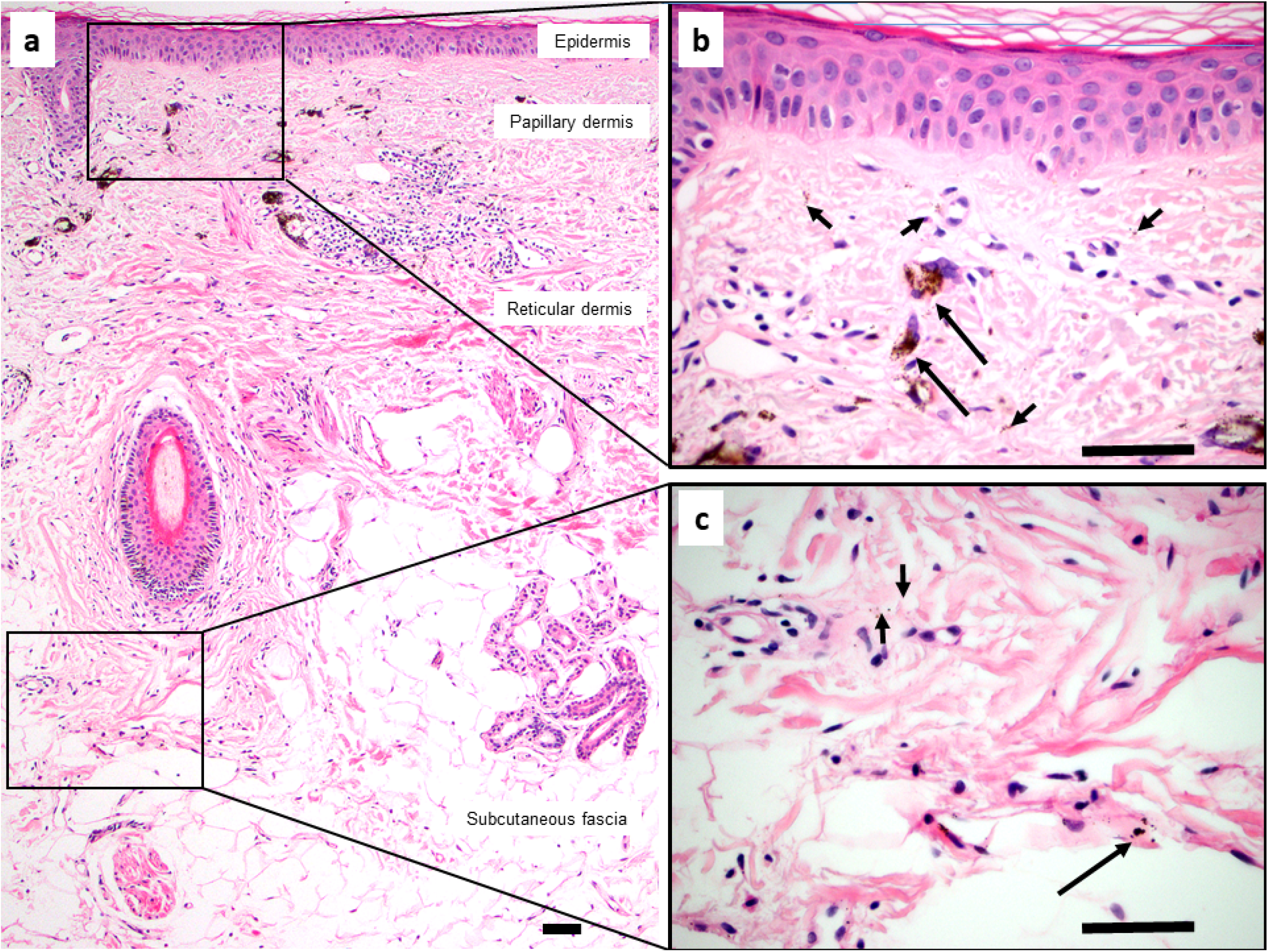
Tattoo pigment in interstitial spaces of the dermis and subcutaneous fascia. **a.** H&E section of skin and subcutaneous fascia with cosmetically injected brown-black tattoo pigment. Pigment particles are present in papillary and reticular dermis and subcutaneous fascia, visible at low magnification. **b,c.** Higher magnification views of the rectangular areas demonstrate both intracellular particles (within macrophages; long arrows) and extracellular particles (within interstitial spaces; short arrows) of the papillary dermis (**b**), reticular dermis and subcutaneous fascia (**c**). Scale bars = 100 μM.

**Fig. 2.**
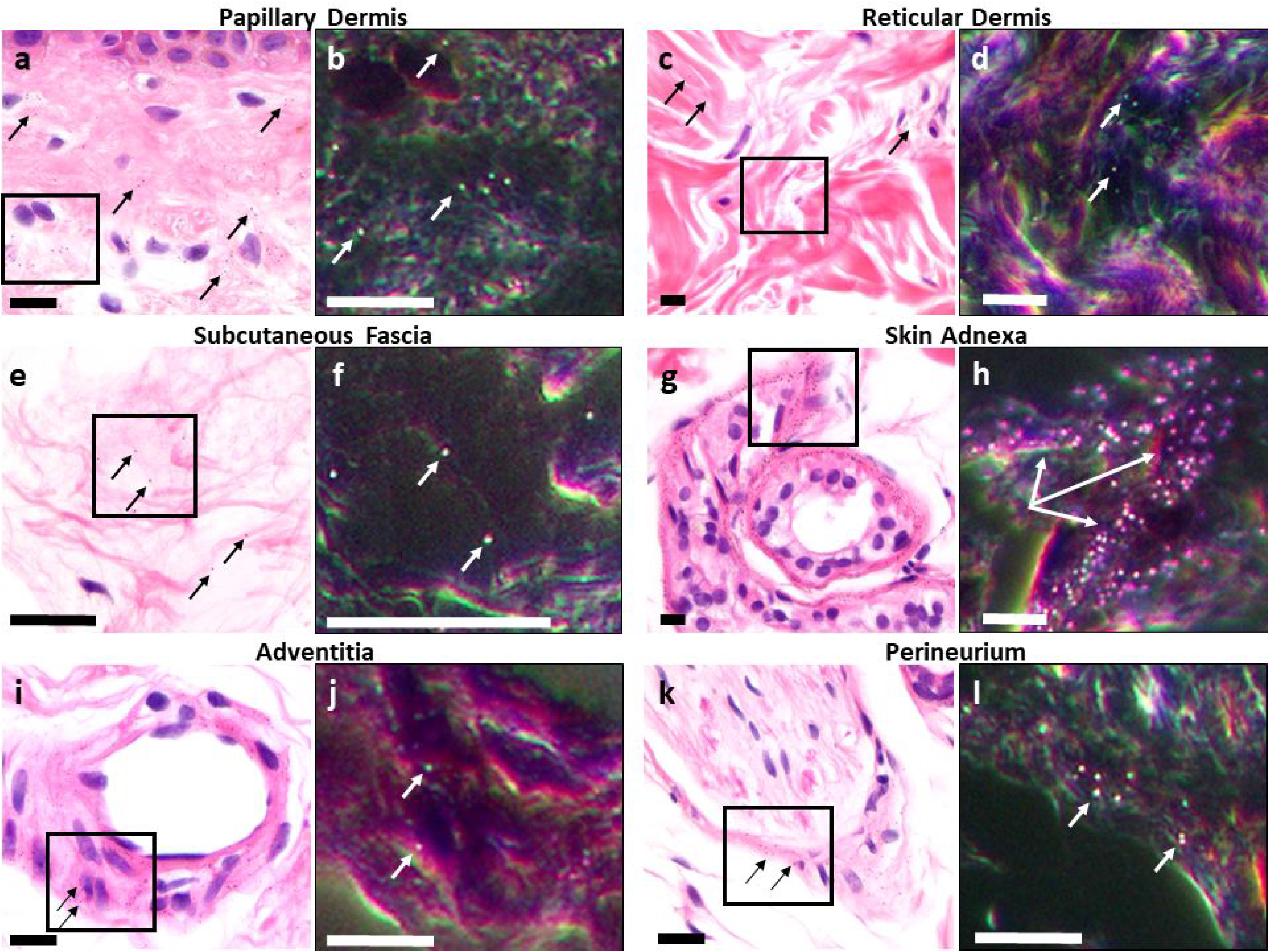
Topically-applied colloidal silver particles are found in the subcutaneous fascia. Juxtaposed light field and dark field microscopy of regions of the same H&E sections: epidermis and papillary dermis (**a,b**), reticular dermis (**c,d**), subcutaneous fascia (**e,f**), adnexa (**g,h**), arteriole with adventitia (**i,j**) and peripheral nerve (**k,l**). Silver particles (which appear as very fine brown-black granules on light microscopy, some labeled with black arrows, and bright granules on dark field microscopy, some labeled with white arrows) are identified in the interstitial spaces of papillary and reticular dermis, and those of the subcutaneous fascia, as well as in the basement membrane of adnexal structures, in perivascular adventitia, and in perineurium. Scale bars = 50 μM.

Colon resection specimens with endoscopically-injected tattoos also demonstrated pigment particles distant from the original submucosal injection site. In samples from all 5 specimens studied, pigment particles were identified not only in colonic submucosa, but also in the muscularis propria and mesenteric fascia (**Fig. 3a**). In a similar fashion to the findings in the skin, pigment particles were both intracellular and within interstitial spaces of the collagenous network of colonic submucosa, muscularis propria and mesenteric fascia (**Fig. 3b-d**). We previously demonstrated such movement of tattoo pigment from colonic submucosa to draining lymph nodes of the mesentery.^11^

**Fig. 3.**
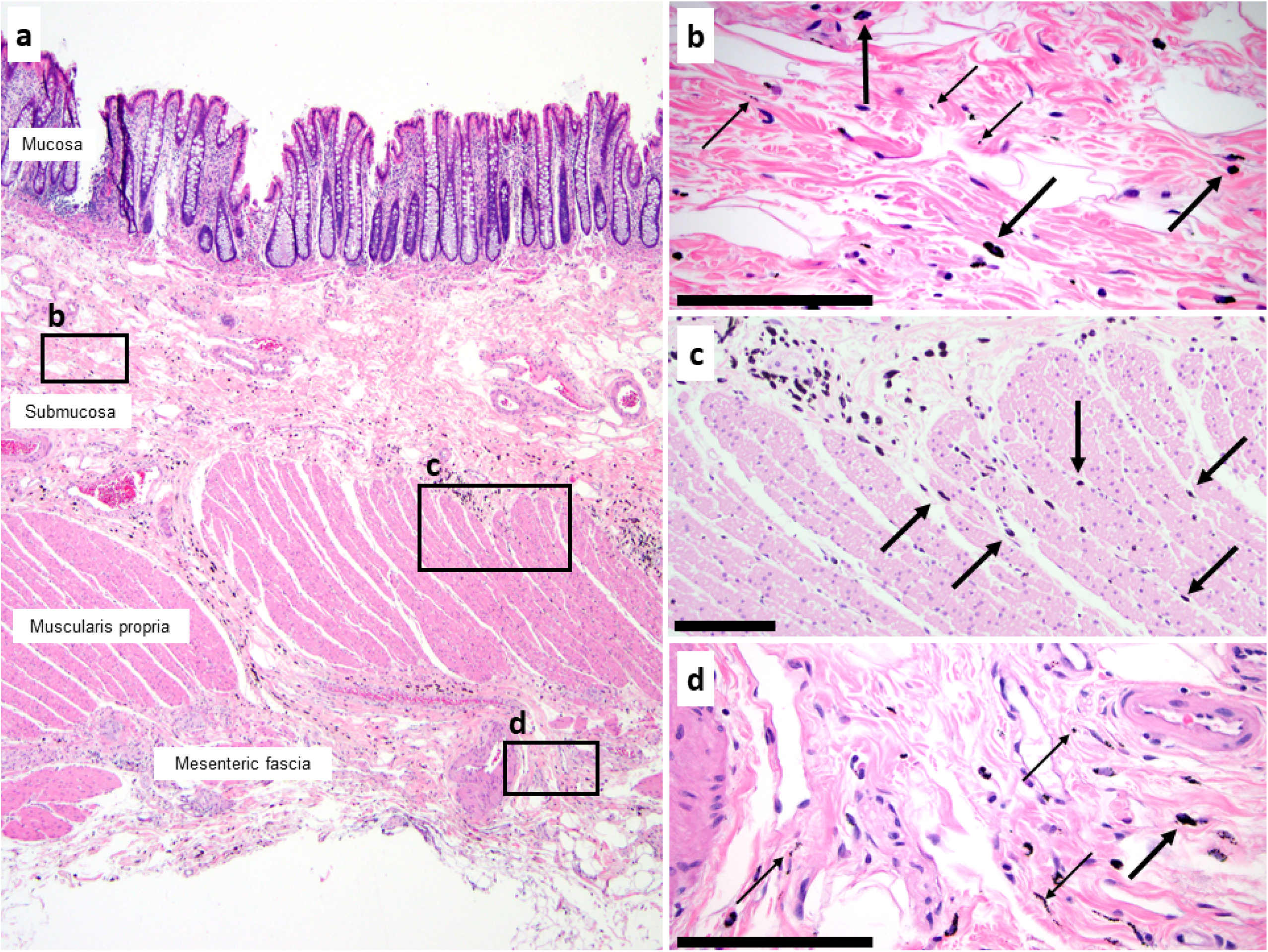
Tattoo pigment in interstitial spaces of the colon and mesenteric fascia. **a.** H&E section of colon resection specimen with endoscopically injected tattoo pigment. Pigment particles are present in submucosa, muscularis propria, and mesenteric fascia. **b.** Higher magnification view of the submucosa demonstrates both intracellular particles (within macrophages; thick arrows) and extracellular particles (within interstitial spaces; thin arrows). **c** Intermediate power view of the muscularis propria shows pigment-containing macrophages (thick arrows) within interstitial spaces between muscle bundles (compare to Fig. 10c for similar display of movement by carcinoma). **d.** Mesenteric fascia has intracellular and intercellular pigment as seen in all other layers. Scale bars = 100 μM.

Because particles are found within macrophages which were shown previously to migrate to regional lymph nodes,^11^ one possible explanation for the appearance of extracellular particles at a distance is that they were carried there intracellularly and then released. Additionally, it is possible with the tattoo pigment that the initial injection was deep enough to explain our findings, though this would not explain the silver particles as they were absorbed from a topical application. We therefore measured the diameter of the extracellular tattoo pigment particles as a function of the depth of their location in the bowel (**Fig. 4a**). Particles in deep mesenteric interstitial spaces were significantly smaller than those in more superficial compartments; the mean particle size in deep mesenteric fascia was 0.46 μm versus 0.61 μm in superficial mesenteric fascia and muscularis propria, and 0.76 μm in submucosa (p<0.01 for all comparisons; **Fig. 4b**). These data suggest that particles were carried via fluid flow rather than via cells, which would have likely resulted in an even distribution of sizes regardless of distance.

**Fig. 4.**
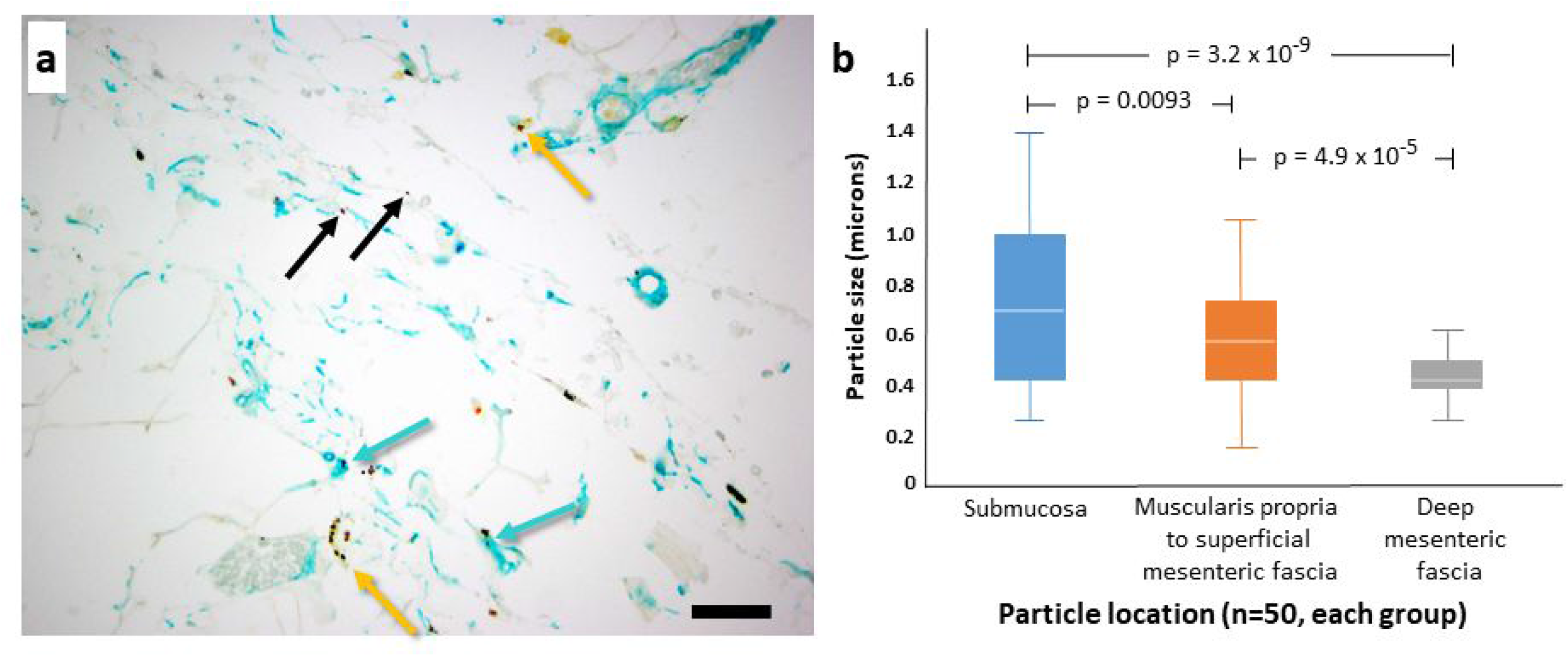
Tattoo pigment particle distribution in the colon. **a** Pericolic mesenteric tissues with tattoo pigment within interstitial spaces (black arrows), within macrophages (yellow arrows), and within interstitial lining cells (teal arrows). Selected areas in colonic submucosa, muscularis propria/superficial mesenteric fascia, and deep mesenteric fascia were examined at high power magnification. Double immunostain for CD68 (macrophage marker; yellow) and CD34 (interstitial lining cell marker; teal). **b** Distribution of particles by size in compartments at increasing distance from lumen. Scale bars = 100 μM.

### Hyaluronic acid staining shows continuity between interstitial spaces across organ boundaries

HA is found in interstitial spaces throughout the body at all stages of development.^10,14–16^ The physical properties of HA suggest that it regulates flow of fluid and other solutes and small molecules within interstitial fluid;^17^ it did not prevent filling by fluorescein in vivo.^11^ We confirmed by staining with HA binding protein (HABP) that it is found broadly in intercellular, pericapillary and perineural, and submucosal and dermal interstitial spaces. This staining showed that non-vascular spaces that appear as white and therefore “empty” by H&E staining are not, in fact, empty, but contain HA (**Fig. 5**). The smallest interstitial spaces between cells are filled with HA including those between epidermal keratinocytes (**Fig. 5a,b**) and within dermal nerve fibers (**Fig. 5g,h**). Pericapillary scale interstitial spaces (lamina propria of colon) were similar (**Fig. 5c,d**). Larger, fibroconnective tissue interstitial spaces filled by HA were observed in dermis (**Fig. 5a,b**) and submucosa and in peri-arterial (**Fig. 5e,f**) and perineurial fibroconnective tissues (**Fig. 5g,h**). HA is thus a surrogate marker of many if not most interstitial spaces. Staining for HA in colon demonstrated that there were connections between all layers from lamina propria through muscularis mucosae to submucosa, and then through muscularis propria into subserosa and mesenteric fascia (**Fig. 6**). Continuity between interstitial spaces in the submucosae and the perivascular adventitia and perineurium in the bowel wall was also evident (**Fig. 7**).

**Fig. 5.**
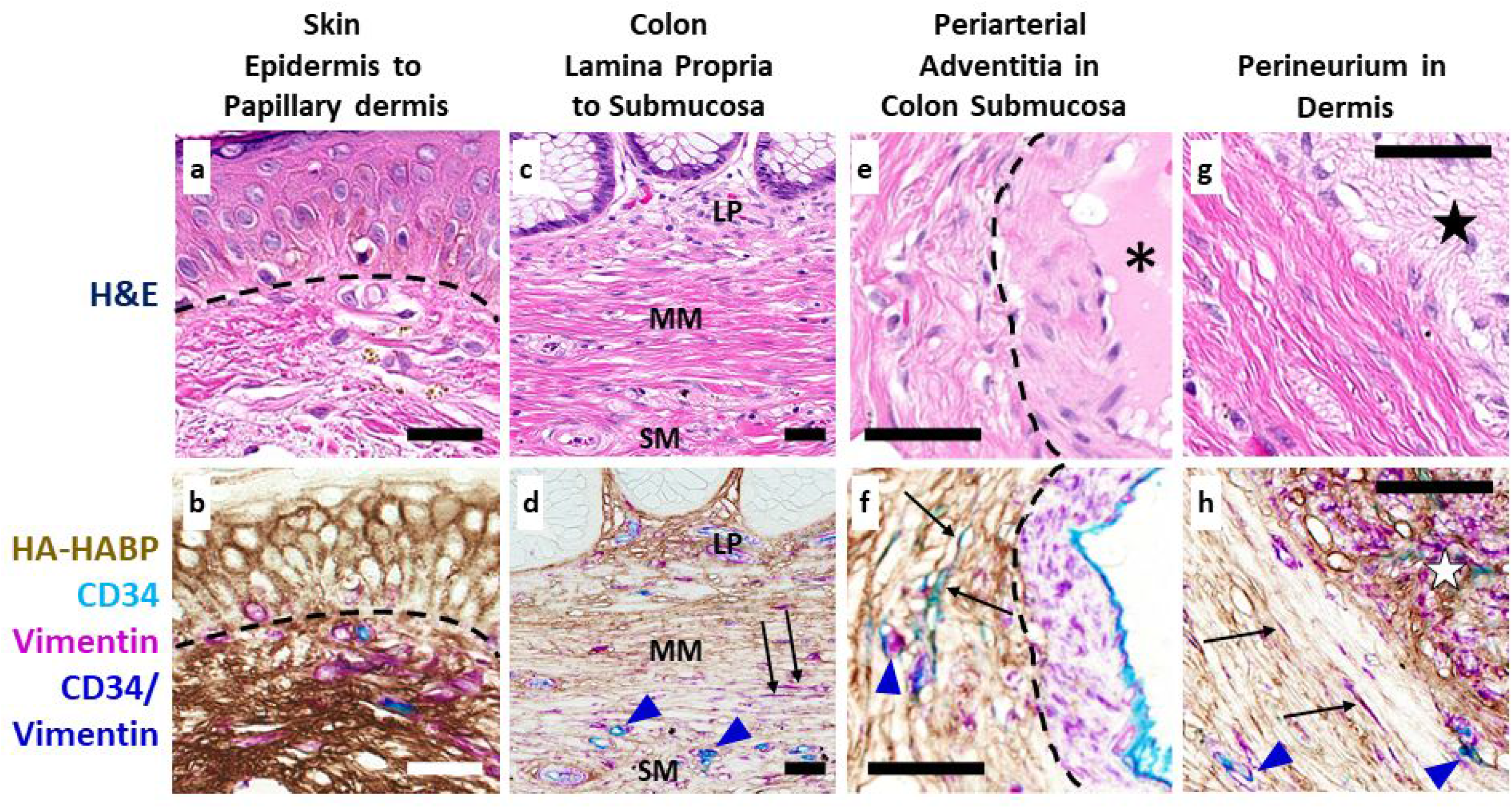
Hyaluronic acid is a marker of interstitial spaces. Full thickness skin and colon samples were stained with H&E (top panels), then decolorized and restained using a multiplex chromogenic assay HABP (brown), vimentin (magenta), and CD34 (teal), CD34/Vimentin overlap (navy blue). **a,b** HA in interstitial spaces between epidermal keratinocytes (above dotted line) and between collagen bundles of the papillary dermis (below dotted line). **c,d** HA in interstitial spaces of the lamina propria (LP) and in channels through the muscularis mucosae (MM) and submucosa (SM). **e,f** HA in interstitial spaces of the adventitia around an artery (lower right, demarcated by dotted line) in the wall of the colon. The lumen of the artery is filled with red blood cells (*, upper panel) and the muscular wall of the artery is highlighted by vimentin staining (magenta; lower panel). **g,h** Interstitial spaces of perineurium and fibroconnective tissue around the nerve (star) in the colonic wall stain for HA, as do the intercellular spaces in the nerve itself. In all multiplex stained images, vimentin staining highlights fibroblasts, mononuclear inflammatory cells, vascular smooth muscle, nerve, vascular endothelial cells, and interstitial lining cells. CD34 stains endothelial cells and interstitial lining cells. Co-localization between CD34 and vimentin (navy blue) indicates either vascular endothelium, when lining capillaries or arteriovenous structures (navy blue arrowheads), or interstitial lining cells when along collagen bundles. Scale bars = 100 μM.

**Fig. 6.**
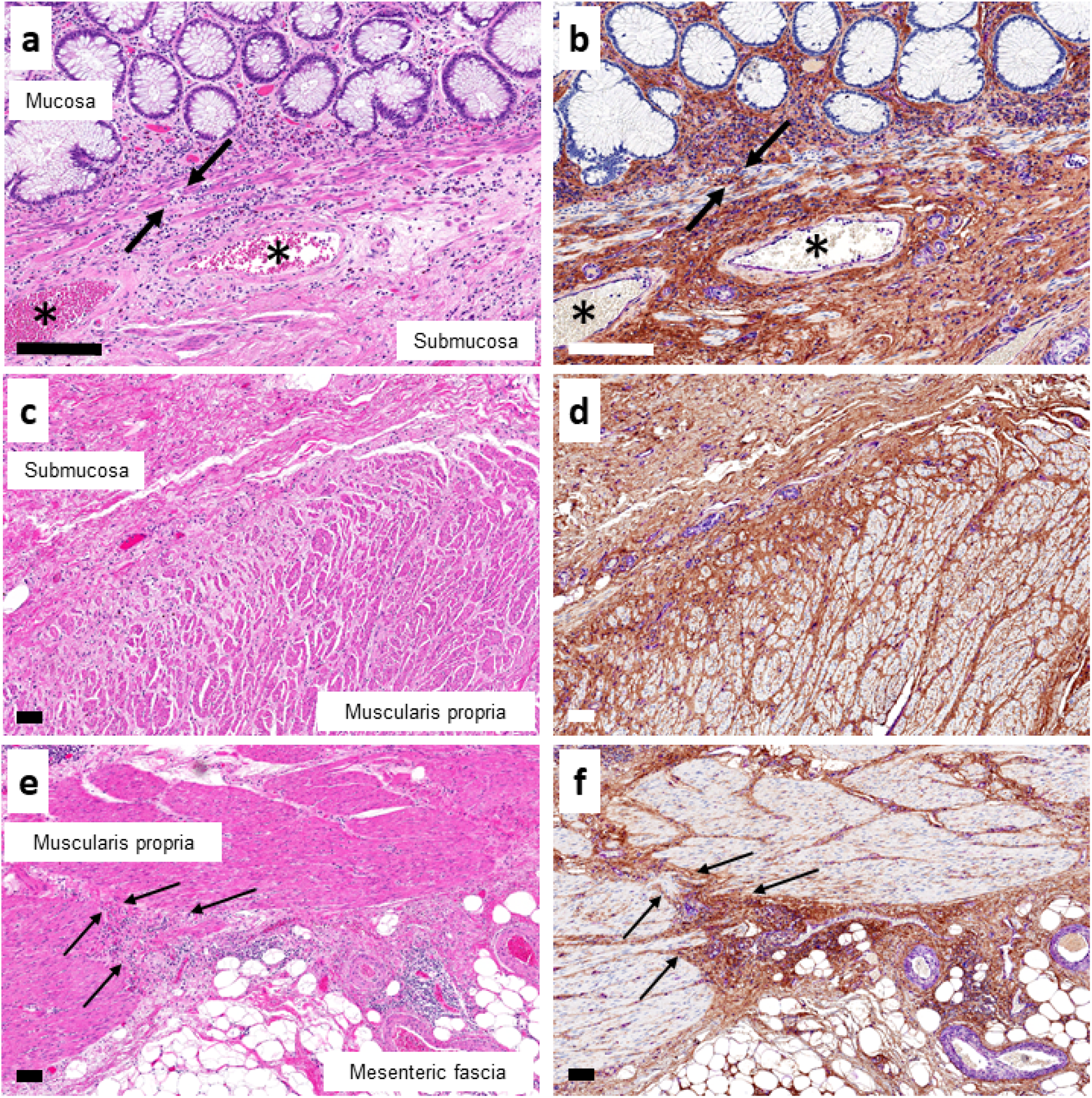
HA staining in interstitial spaces is continuous through all the layers of the colon into mesenteric soft tissues. Adjacent colon samples were stained with H&E (left) and a triplex chromogenic assay (right) for HABP (brown), vimentin (magenta), and CD34 (teal). **a,b** Juncture of lamina propria to submucosa, with channels observed through the muscularis mucosae (black arrows). There is continuity between all these layers by HA staining (right). * marks large veins in submucosa with HA staining showing continuity between perivascular stroma and the surrounding submucosal stroma. (40X magnification). **c,d** Juncture between colonic submucosa and the muscularis propria. HA staining shows continuity of HA-filled interstitial spaces from submucosa through small pericapillary channels between muscle bundles of the muscularis propria (right). (10X magnification). **e,f** Edge of fibrovascular bundle (arrows) passing between colonic wall, through muscularis propria, into subserosa and mesenteric fascia (right). In all multiplex-stained images, staining is as in Fig. 5. Scale bars = 100 μM.

**Fig. 7.**
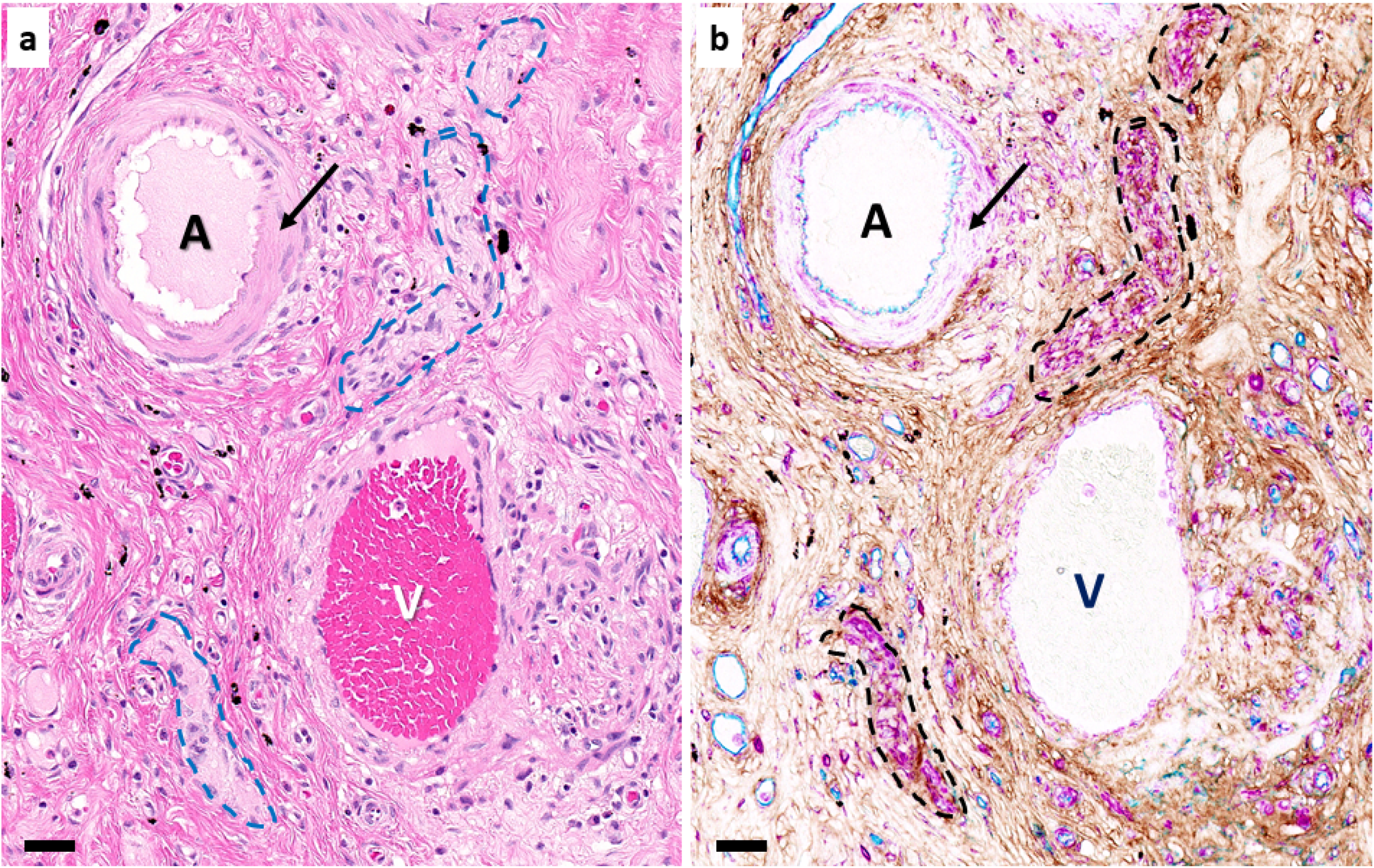
Continuity between interstitial spaces, perivascular adventitia and perineurium in colonic submucosa. **a** H&E section of colonic submucosa with artery (A), vein (V), and peripheral nerve (dashed lines) Arterial tunica media (muscular coat) labeled by arrow. **b** HABP-vimentin-CD34 triplex chromogenic stain of the same tissue section after de-colorization (HABP, brown; vimentin, magenta; CD34, teal). Nerves show strong vimentin staining as does the tunica media (of the artery (arrow). HABP highlights HA in interstitial spaces, showing continuity of spaces between connective tissue of the bowel and the perivascular and perineurial stroma. A, artery; V, vein. In all multiplex-stained images, staining is as in Fig. 5. Scale bars = 100 μM.

In the skin, HA-filled interstitial spaces were continuous from intercellular spaces of the epidermis into papillary dermis (**Fig. 5a,b, 8a,b**) and then to the reticular dermis (**Fig 8a-d**) and deeper into subcutaneous fascia (**Fig. 8a,b, e,f**). Intercellular and pericapillary interstitial spaces of subcutaneous adipocytes are continuous with the interwoven subcutaneous fascia and with perivascular adventitia (**Fig. 8e,f**).

**Fig 8.**
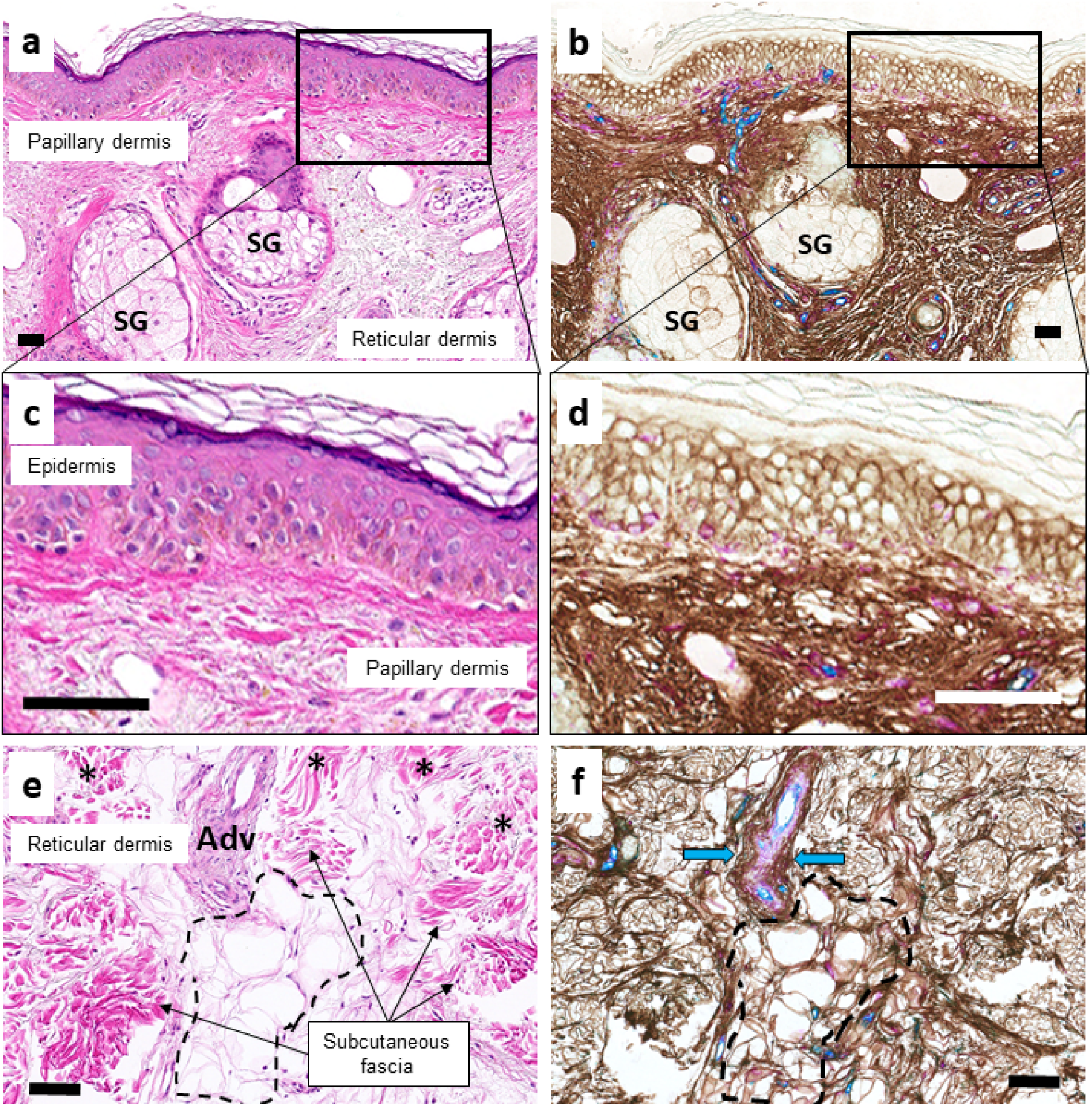
Continuity of interstitial spaces through all layers of the skin into subcutaneous fascia. Adjacent colon samples were stained with H&E (left) and a triplex chromogenic assay (right) for HABP (brown), vimentin (magenta), and CD34 (teal). **a,b** Junctional region between papillary dermis and reticular dermis with adnexal sebaceous glands (SG). HABP staining of HA shows continuity from epidermal interstitial spaces through to the reticular dermis. (10X magnification). **c,d** Rectangular regions from the top images expanded showing continuity from epidermal intercellular spaces into the papillary dermis (60X magnification). **e,f** The lower reaches of papillary dermis (*) extending into the subcutaneous soft tissues with peri-vascular adventitia (Adv) and subcutaneous fascia separated by adipose tissue. HABP staining highlights continuity between all these spaces including intercellular spaces and pericapillary spaces between adipocytes (yellow dashed line) and the larger spaces of the reticular dermis, subcutaneous fascia and perivascular adventitia (blue arrows). In all multiplex-stained images, staining is as in Fig. 5. Scale bars = 100 μM.

Similarly, the space of Mall in the portal tracts of the liver shows HA staining between the interstitial spaces of stroma around the intrahepatic bile duct and of the stroma around the hepatic artery and portal vein, indicating continuity (**Fig. 9**).

**Figure 9.**
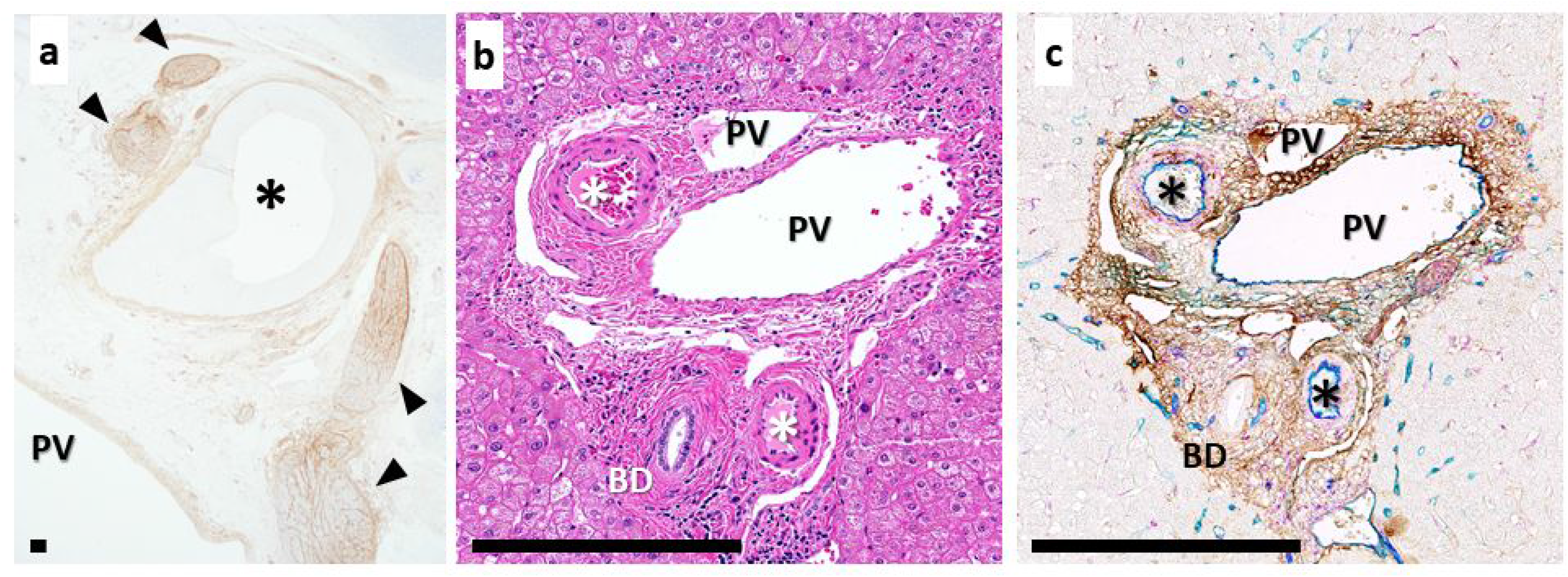
Continuity of extrahepatic interstitial spaces of the porta hepatis and intrahepatic spaces within the “space of Mall” demonstrated by HA localization. **a** HA staining of interstitial spaces of structures of the portal hepatis: the adventitia of portal vein (PV), hepatic artery (*) and within and around nerves (arrowheads). (Single stain of HA by HABP binding, DAB). **b** H&E section of liver with portal triad: hepatic artery (*), portal vein (PV), and bile duct (BD). **c** Multiplex chromogenic assay of the same region. HABP (brown), vimentin (magenta), CD34 (teal). HABP staining is coincident with the space of Mall; it highlights HA in interstitial spaces, showing continuity between all compartments of the space of Mall. In all multiplex stained images, staining is as in Fig. 5; interstitial lining cells are in the space of Mall. Scale bars = 400 μM.

### Multiple tumor types move through interstitial spaces

Movement of tumor through these interconnected interstitial spaces is demonstrated in cases of peribiliary spread of cholangiocarcinoma, which tracks between collagen fibers through the biliary submucosa and throughout the space of Mall, while respecting the rigid boundary of the limiting plate of hepatocytes (**Fig. 10a, b)**.^18,19^ Similarly, colon adenocarcinoma may be seen moving between collagen bundles and between muscle bundles of the muscularis propria into the mesentery (**Fig. 10c,d**) and malignant melanoma is known to give rise to “in transit” metastases within the dermis without evidence of lymphovascular invasion (**Fig. 10e**).^20,21^ The presence of melanoma tumor nests between collagen bundles (**Fig. 10f**) without an associated desmoplastic reaction, similar to the appearance of macrophages with tattoo pigment in Fig 3c, supports that the cells are within pre-existing interstitial spaces and have not elaborated tissue-destructive digestive enzymes.

**Figure 10.**
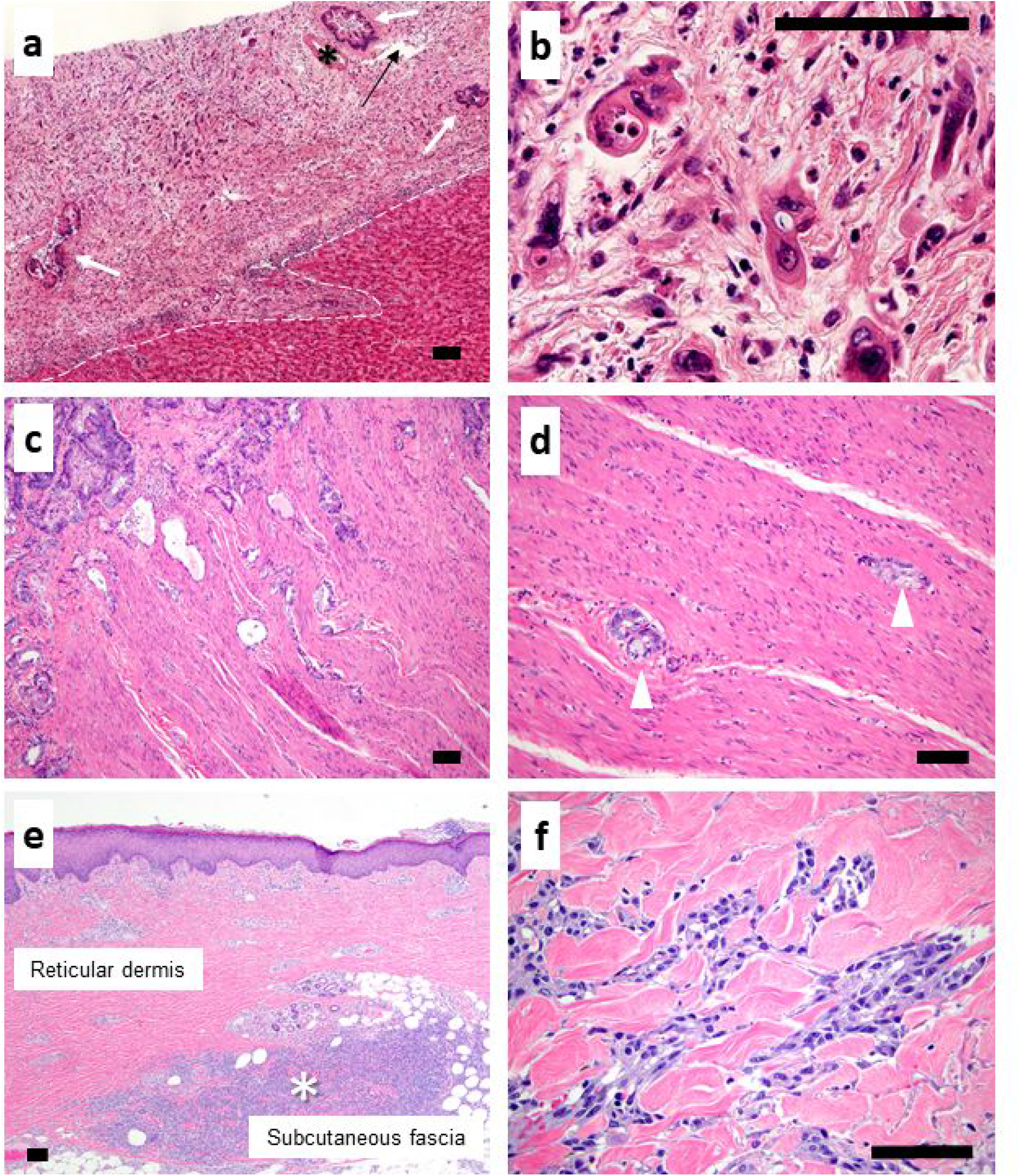
Interstitial spaces are a route of spread for malignant tumors. **a** Periductal spread of hilar cholangiocarcinoma through the space of Mall of the portal tract stroma in the liver. Note the absence of invasion through the periportal limiting plate (dotted line) with tumor confined completely within the portal tract stroma, surrounding many portal structures, but not invading them, i.e. bile ducts (white arrows), hepatic artery (*) and portal vein (black arrow). **b** Cholangiocarcinoma within interstitial spaces of portal tract stroma; same tumor as **a**. **c** Colonic adenocarcinoma (dark blue/purple glands percolating from upper left, downward) infiltrating through the submucosa and between muscle bundles of the muscularis propria. **d** Same tumor as **c** showing tumor nests (arrow heads) between pink muscle bundles without desmoplastic reaction. **e** In-transit malignant melanoma tumor deposit (*) in deep reticular dermis and subcutaneous fascia, several centimeters away from the primary lesion. **f** Higher magnification view of melanoma (grey and blue cells) in **e** infiltrating through interstitial spaces of reticular dermis between its pre-existing, pink, acellular collagen bundles. All images H&E. Scale bars = 100 μM.

## Discussion

We demonstrate here that non-biological pigment particles (cosmetic tattoos and colloidal silver in skin and endoscopically-injected tattoo pigment in colon) are found across classically defined tissue boundaries from their sites of entry into the body, within subcutis and mesentery, respectively, indicating movement across compartments traditionally thought to be anatomically separate. We also use HABP staining to show that interstitial spaces are continuous between tissue compartments and fascial planes in the colon, skin, and liver, as well as within the fibrous tissues around blood vessels and nerves. Taken together, the data indicate that there is continuity of these interstitial spaces within organs (shown here for skin and colon) and outside of organs (shown here for skin into subcutis and colon into mesentery).

Although movement of pigment particles could occur via macrophage engulfment and migration, our data suggest that this is unlikely. The significantly smaller mean particle size in progressively deeper layers of the bowel wall suggests that particles were mostly carried via fluid flow rather than by macrophage carriage and subsequent release, as cell-mediated transport would have resulted in an even distribution of sizes regardless of distance. Absence of a need for cell-based particle movement is similarly highlighted by the presence of colloidal silver through all layers of skin, without evidence of macrophage engulfment (**Fig. 2)**. The argyria specimens also make it unlikely that the spread of tattoo pigment injection was due to mechanical pressure from the injection: the colloidal silver was applied as a topical agent on the skin, diffusing through the epidermis into the deeper layers without instrumentation. Its dissemination into sub-epidermal stroma and then into the epithelia of the adnexal structures suggests that basement membranes are permeable to movement of minute extracellular particles (**Fig. 2)**.

The large interstitial spaces, which appear as empty white spaces on routine formalin-fixed, paraffin-embedded histochemically-stained specimens, are filled with HA. This finding allowed us to demonstrate that interstitial spaces are continuous between tissue compartments and fascial planes in the colon, skin, and liver, as well as within the fibrous tissues around blood vessels and nerves, which may pass through multiple organs.

While this has not, to our knowledge, been demonstrated previously for skin and colon and their adjacent tissues, it confirms Mall’s experimental demonstration of continuity within the portal tract stroma.^1^ Mall demonstrated that pigmented gelatinous substances (cinnabar and Prussian blue) distributed into the microanatomic structures of the portal tract following their injection into the vascular supply of cat livers. Based on Mall’s experimental observations, the perivascular and periductal portal tract stroma can be visualized as a unified network of fibers with intervening spaces that communicate with each other and are in continuity with the vascular (entrance) and lymphatic (egress) systems in the liver. Our findings are in agreement with Mall’s observations: HA staining was present in a continuous fashion within portal tract stroma ensheathing hepatic arteries, portal veins, and the biliary tree, in support of there being a continuous space. The intrahepatic portal triads show how spaces that are in and around structures including arteries, veins, nerves, and ducts (such as ureters, urethra, epididymis, Fallopian tubes, salivary gland exocrine ducts) that pass through multiple tissues and organs could serve as conduits for interstitial fluid.

Particle movement by flow through fluid-filled channels likely arises from external and internal (physiologic) mechanical forces such as peristalsis of the gut, positional or mechanical pressure on the skin and subcutaneous tissues, and rhythmic compression of perivascular/adventitial stroma resulting from arterial wall expansion in systole. Diaphragmatic movement between inhalation and exhalation will also cause rhythmically oscillating pressures on tissues in the thorax and the abdomen that may not have strong driving pressures from other sources, such as the mesentery. Movement of fluid through interstitial spaces of the perineurium may be subject to all these influences, as well as limited by the phenomenon of interstitial exclusion.^17^

Conventional models suggest that the extracellular matrix serves as a physical barrier to cancer cell migration and that destructive breakdown by matrix metalloproteinases is a prerequisite step for cancer cell invasion and metastasis. However, extensive work has shown that cancer invasion, at least in its initial steps, is largely non-destructive, without significant tissue remodeling, and that malignant cells can traffic through pre-existing interstitial spaces, which likely serve as routes of least resistance.^22–26^ In keeping with this work, our findings, as shown in **Fig. 10**, may explain some features of classically recognized cancer behavior such as tumor cell spread within an organ tissue plane (single-cell filing of lobular carcinoma of the breast, linitis plastica of the stomach and, as shown in this study, periductal spread of cholangiocarcinoma). Likewise, continuity across tissue planes could explain the clinically recognized “discontinuous” spread of cancer such as mesenteric tumor deposits of colorectal cancer and subcutaneous melanoma “in transit” metastases (**Fig. 10c-f**).

These findings have important implications for other processes such as infections, autoimmunity, and host-microbiome interactions. Interstitial spaces across the human body could act as pathways for both commensal micro-organisms of the human microbiome and various pathogens. Direct continuity across tissue planes may explain necrotizing fasciitis, a fulminant form of soft tissue infection resulting from virulent bacterial strains that gain access to the interstitial spaces and cause widespread necrosis of subcutaneous and perimuscular fascia and even cross the blood:brain barrier to cause meningitis.^27^ Continuity across the layers of the intestine and through the mesenteric and portal vein adventitia may provide a direct route for the translocation of gut bacteria to the liver (“gut-liver axis”).^28,29^ The existence of these pathways, in parallel with the mesenteric lymphatic and portal venous systems, may play a significant role in gut microbiota signaling not only in chronic liver disease and cirrhosis, but also in the context of autoimmune disorders, both liver-specific (autoimmune hepatitis) and systemic.^30^ New evidence shows that the central nervous system has a system of lymphatic vessels that line the dural sinuses and connect the cerebrospinal fluid space to the cervical lymph nodes; one anatomic route of this connection could be via the peri-arterial adventitia of the carotid arteries.^31,32^ The presence of interconnected interstitial spaces of channels that track along the perineurium of the peripheral nerves provides a potential novel route of communication between the gut and the brain (“gut-brain axis”).^33^

We demonstrate that the fibrous layers of nerves and blood vessels have no discrete separation from the fibrous components of the organs through which they travel and that, likewise, their interstitial spaces are not segregated from each other. Thus, we suggest that there is continuity of fibrous tissue interstitial spaces within and between organs and that these spaces are also potentially continuous between more distant parts of the body, at least along the vasculature and nerves. We speculate that these spaces serve as pathways for molecular signaling and cell trafficking in a way that is both in series and in parallel with the established pathways of the cardiovascular and lymphatic systems, although flow is likely limited by the structural proteins and glycosaminoglycans of the interstitium.^17^ Given that the estimated volume of interstitial fluid in the body is more than 3 times the combined fluid volume of the cardiovascular and lymphatic systems,^34^ the existence of an interconnected interstitial compartment suggests an anatomic basis for understanding both physiological processes and disease pathophysiology.

## Methods

### Patients and Tissue Specimens

Formalin-fixed, paraffin-embedded (FFPE) archival anatomic pathology tissue blocks were collected from a) surgical segmental colectomy specimens performed for the treatment of malignant polyps (5 patients), all of which contained tattoo pigment (India ink) that was injected into colonic submucosa adjacent to the lesion at the time of colonoscopy and prior to surgery, b) skin punch biopsy specimens which included cosmetic tattoos (3 patients), c) two skin punch biopsy specimens containing colloidal silver in a patient with argyria after topical skin application of colloidal silver (1 patient), and d) resection specimens of otherwise normal liver containing metastatic tumor (4 patients). Inclusion criteria were: bowel specimens – the presence of all anatomic compartments (mucosa, submucosa, subserosa and mesentery) within the tissue section; skin specimens – the presence of epidermis, dermis, and subcutaneous fascia and adipose tissue. Samples from patients less than 18 years old were excluded. The study was conducted in accordance with the guidelines and regulations and with the approval of the New York University Langone Health Institutional Review Board.

### H&E Staining, Scanning and Decolorization

FFPE specimens were sectioned at 5 μm onto charged slides (Fisher Scientific, Cat # 22-042-924). Slides were dried for 1 h at 60°C, deparaffinized in xylene, rehydrated through a graded series of ethanols, and rinsed in distilled water. Slides were hematoxylin (Richard-Allan Scientific, Cat# 7211) and eosin (Leica, Cat# 3801619) stained using standard laboratory protocol.^35^ Upon completion of staining, slides were dehydrated through a series of ethanols and xylene, and mounted with Cytoseal 60 (Richard-Allan Scientific, Cat# 8310-4). Slides were scanned using a Leica Biosystems Aperio AT2 System and digitally archived via eSlide Manager (Version 12.3.2.5030). Following scanning, H&E slides were immersed in xylene to remove coverslips. Slides were rehydrated through xylene, graded ethanol, and running distilled water, decolorized with 10% acetic acid in 70% ethanol for 1 h, rinsed in distilled water, and then further decolorized in 70% ethanol for 2-3 h. Evaluation for the decolorization end-point was checked every 30 min.^36,37^ Once decolorization was complete, slides were rinsed in running distilled water.

### Multiplex Immunohistochemistry

Unconjugated murine anti-human Vimentin (Ventana Medical Systems, Cat# 790-2917, RRID: AB_2335925) clone V9, unconjugated murine anti-human CD34 (Ventana Medical Systems, Cat# 790-2927, RRID: AB_2336013) clone QBEnd/10, and unconjugated murine anti-human CD68 (Ventana Medical Systems, Cat# 790-2931, RRID: AB_2335972) clone KP1 were used for chromogenic immunohistochemistry. Biotinylated HABP (Calbiochem, Cat# 385911) was used for a non-immune chromogenic assay. The protein binds specifically to HA (≥ 2000 M.W).^38^ Biotinylated HABP is directly detected using a streptavidin peroxidase-DAB detection system.

Chromogenic immunohistochemical multiplexing (mIHC) was performed on a Ventana Medical Systems Discovery Ultra using Ventana reagents except as noted, according to the manufacturer’s instructions and best practices.^39,40^ Slides were dried in a 60°C incubator for 1 h and deparaffinized on-instrument. Decolorized samples for mIHC bypass prerun incubation and deparaffinization and are started directly from buffer. All sample sets were run with a tissue microarray as a positive, negative and mIHC crossover controls.

For the HABP-Vimentin-CD34 triplex assay, endogenous peroxidase was blocked with 3% hydrogen peroxide for 8 min at 37°C. HABP was applied at 0.5μg/ml (1:100 dilution) in tris-buffered saline with 1% bovine serum albumin (TBSA) and incubated for 12 h at room temperature. The biotinylated protein was then directly detected using horseradish peroxidase-conjugated streptavidin with DAB substrate. Sections were then antigen retrieved using Cell Conditioner 1 (Tris-Borate-EDTA ph8.5) for 20 min at 95°C. Peroxide blocker was reapplied as above. Vimentin antibody was applied without dilution and incubated for 20 min at 37°C followed by a goat anti-mouse horseradish peroxidase (HRP)-conjugated multimer applied for 8 min at 37°C. This was detected with purple (tyramide-TAMRA) chromogen for 8 min at 37°C. Subsequently, sections were denatured in Reaction Buffer (Cat# 950-300) for 32 min at 95°C. Endogenous peroxidase was blocked as previously described. CD34 was applied neat for 60 min at 37°C. A goat anti-mouse secondary HRP-conjugated multimer was applied neat for 8 min at 37°C and detected using teal (tyramide-Cy5) chromogen for 16 min at 37°C. Slides were subsequently dehydrated, cover-slipped and scanned.

Extracellular versus intracellular localization of pigment particles was assessed on sections stained with duplex immunohistochemistry for CD68 (macrophage marker) and CD34 (marker of interstitial lining cells). For the CD68-CD34 duplex assay, CD34 antibody was applied neat to deparaffinized slides for 60 min at 37°C. Goat anti-mouse secondary HRP-conjugated multimer secondary was applied neat for 8 min at 37°C and detected using Teal chromogen for 8 min at 37°C. Slides were denatured in reaction buffer for 32 min at 95°C followed by antigen retrieval using Cell Conditioner 1 for 20 min at 91°C. CD68 was applied neat for 32 min at 37°C. A goat anti-mouse secondary alkaline phosphatase-conjugated multimer was applied for 8 min at 37°C and detection was completed using Yellow (tyramide-dabsyl) chromomgen for 8 min at 37°C. Slides were subsequently dehydrated, coverslipped and scanned.

#### Particle Size Measurement and Statistical Analysis

Sizes of 50 tattoo pigment particles were measured using eSlide Manager digital annotation ruler tool. It was not possible to measure the size of particles within macrophages because particles were often stacked up on each other in the cytoplasm. Student t test was used to analyze discrete data. All statistical analyses were two-tailed. P values less than 0.05 were considered statistically significant.

## Acknowledgments

The NYULH Center for Biospecimen Research and Development, Histology and Immunohistochemistry Laboratory (RRID:SCR_018304) is supported in part by the Laura and Isaac Perlmutter Cancer Center Support Grant: NIH/NCI P30CA016087 and the National Institutes of Health S10 Grants; NIH/ORIP S10OD01058 and S10018338.

